# Multiplex PCR based Detection Methods of Common Plant Transgenes

**DOI:** 10.64898/2026.04.23.720246

**Authors:** Atsuko Iuchi, Satoshi Iuchi, Yukie Aso, Hiroshi Abe, Masatomo Kobayashi, Taiji Kawakatsu

**Author notes:** **Corresponding author** Taiji Kawakatsu. These authors contributed equally to this work (Atsuko Iuchi, Satoshi Iuchi).

## Abstract

Accurate verification of transgenic plant materials is essential for maintaining scientific integrity and ensuring experimental reproducibility. As the number and diversity of transgenic constructs continue to expand, there is a growing need for practical and scalable methods that enable routine confirmation of transgene presence and identity. Reliable detection systems are particularly important for laboratories handling large numbers of genetically modified lines or distributing materials across research groups. To address this need, we developed two complementary methods for efficient detection of commonly used transgenes. The first method, fDET, is a higher-throughput system capable of simultaneously detecting 15 transgenes and three endogenous genes in a single multiplex PCR reaction followed by capillary electrophoresis. This approach provides rapid, high-resolution detection suitable for high-volume or time-sensitive applications. The second method, DET, offers a more accessible workflow that detects 10 transgenes and one endogenous gene using four multiplex PCR reactions followed by agarose gel electrophoresis. Because DET requires only standard molecular biology equipment, it can be readily implemented in a wide range of laboratory environments without specialized instrumentation. Together, these methods provide flexible and practical solutions for verifying the genetic status of both transgenic and non-transgenic plant materials. By enabling efficient and comprehensive transgene detection, they support reproducible experimentation, facilitate quality control in plant research, and streamline the management and exchange of genetically modified lines. These approaches contribute to more reliable and transparent use of transgenic resources across the plant science community.

## Introduction

Genetic transformation is a fundamental technique in plant biology and biotechnology, enabling functional gene analysis and the introduction of novel traits into plants and crops. Accurate identification of transgenes is therefore essential for scientific rigor, reproducibility, and transparent resource distribution. The rapid adoption of genome editing has further complicated resource management, as transgenic plants, genome-edited lines, null segregants, and non-transgenic plants are often handled together. Incomplete segregation or insufficient verification can lead to uncertainty in genetic status and unintended distribution of transgenic material. Efficient detection of commonly used transgenes is thus critical for genetic verification, biosafety compliance, and reliable sharing of plant resources to ensure research reproducibility. Multiplex PCR combined with capillary electrophoresis is widely used in plant research because of its high sensitivity and throughput, and has also been applied to transgene detection. Recent studies have demonstrated multiplex fluorescent PCR assays capable of detecting dozens of transgenes, mainly targeting trait-associated elements introduced into crop plants for agricultural improvement (Yi et al. 2022). However, such approaches are generally designed for crop- or trait-specific purposes. In contrast, routine research laboratories and stock centers require universal, efficient screening methods targeting widely used selectable markers and reporter genes shared across diverse species and transformation systems, for which optimized high-throughput frameworks remain limited. To address this need, we developed two PCR-based methods for detecting common plant transgenes. Fluorescent detection of transgenes (fDET) enables high-throughput analysis using a single PCR reaction followed by capillary electrophoresis, whereas detection of transgenes (DET) provides a lower-throughput alternative using conventional PCR and agarose gel electrophoresis.

## Results and Discussion

We selected a total of 15 commonly used transgenes and 3 endogenous genes as detection targets (Figure 1a, Supplemental Table S1). These include four selection marker genes: *BAR* (*BIALAPHOS RESISTANCE*, conferring resistance to glufosinate ammonium, such as BASTA and bialaphos), *HPT* (*HYGROMYCIN PHOSPHOTRANSFERASE*, conferring resistance to hygromycin B), *NPTII* (*NEOMYCIN PHOSPHOTRANSFERASE II*, conferring resistance to kanamycin and related aminoglycoside antibiotics), and *BLA* (***B****ETA-LACTAMASE*, which confers resistance to ampicillin); three constitutive promoters and their associated terminators: Cauliflower mosaic virus 35S, *NOS* (*NOPALINE SYNTHASE*), and *MAS* (*MANNOPINE SYNTHASE*); the *Ac* (*Activator* transposase) gene and its endogenous genomic insertion site for monitoring transposon tagging; three reporter genes: *GFP* (*GREEN FLUORESCENT PROTEIN*), *GUS* (***B****ETA-GLUCURONIDASE*), and *LUC* (*LUCIFERASE*); and *Cas9* (*CRISPR-ASSOCIATED PROTEIN 9*), a widely used genome editing enzyme. Additionally, we included *rbcL* (*RIBULOSE-1,5-BISPHOSPHATE CARBOXYLASE/OXYGENASE LARGE SUBUNIT*) and *CBP20* (*CAP-BINDING PROTEIN 20*) as positive controls, representing chloroplast- and nuclear-encoded reference loci, respectively. The selection marker genes and constitutive promoters/terminators included in our panel are commonly found in the T-DNA regions of major binary vectors used in Arabidopsis transformation, such as pROK2 (SALK lines; Alonso et al. 2003), pDAP101 (SAIL lines; McElver et al. 2001), and pCGN derivatives (Ds lines; Fedoroff and Smith. 1993). We also included *Ac* and its insertion site as detection targets. Although *Ac* is theoretically eliminated from D_s_ lines through negative selection, its unintended retention can enable further transposition of *D*_*s*_ elements. Therefore, we monitor both *Ac* and its insertion site to ensure the integrity and stability of transposon-tagged resources. If *Ac* is present in a homozygous manner, the genomic region harboring the insertion site cannot be amplified. The primer-binding regions of *rbcL* are highly conserved among plant species, making *rbcL* a universal positive control. In contrast, the primer-binding regions of *CBP20* are not conserved across species or even among Arabidopsis accessions, making *CBP20* a positive control specific to major Arabidopsis accessions, such as Col-0 and L*er*. Some functional transgenes (e.g. *Cas9*) may be codon optimized for the host species. In such case, primer sequences should be modified to match the actual transgene sequences.

**Figure 1.**
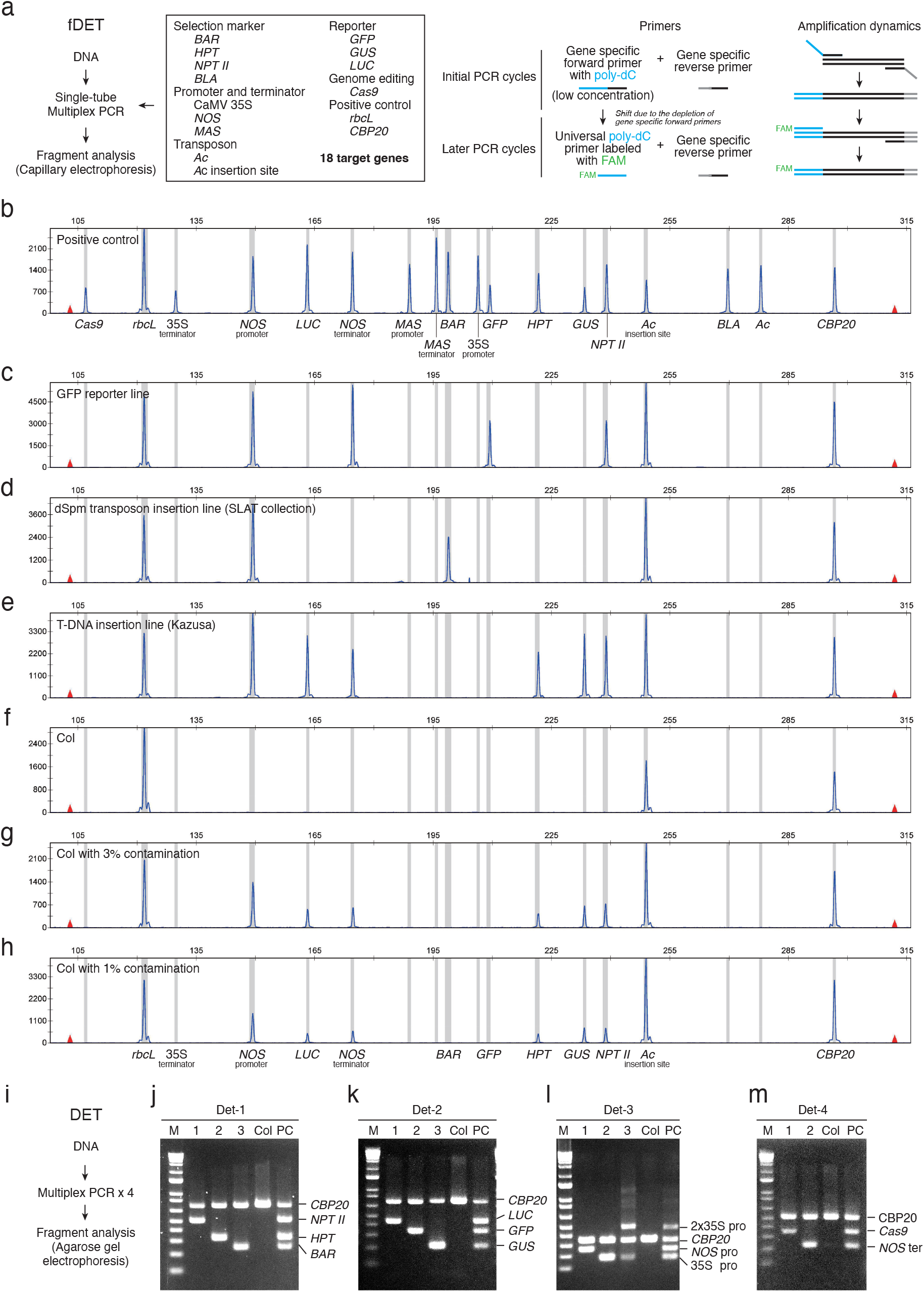
Multiplex PCR based Detection Methods of Common Plant Transgenes. (a) Schematic representation of fDET. A single-tube multiplex PCR combined with capillary electrophoresis–based fragment analysis enables simultaneous detection of up to 18 target genes. During early PCR cycles, gene-specific forward primers with a poly-dC extension add a poly-dC sequence to the 5′ end of target amplicons. After depletion of the low-concentration forward primers, a FAM-labeled poly-dC primer and gene-specific reverse primers amplify the tailed fragments, resulting in fluorescently labeled amplicons. (b) Optimized fDET result using a positive control consisting of pooled genomic DNA extracted from plants harboring various transgenes. (c)-(e) Examples of identification of transgenes by fDET. (c) GFP reporter line, (d) dSpm transposon insertion line from the SLAT collection, and (e) Kazusa T-DNA insertion line. (f)-(h) Sensitivity of fDET. fDET detect contamination as low as 1%. (f) Col (non-transgenic), (g) Col spiked with 3% T-DNA insertion line, (h) Col spiked with 1% T-DNA insertion line. The spiked line is identical to that shown in (e). (i) Schematic representation of DET. Four multiplex PCR reactions followed by agarose gel electrophoresis–based fragment analysis detect up to 11 target genes. (j)-(M) Examples of transgene identification by DET. (J) Det-1 primer mix (*NPTII, HPT* and *BAR*), (K) Det-2 primer mix (*GFP, GUS* and *LUC*), (L) Det-3 primer mix (35S promoter and *NOS* promoter), and (m) Det-4 primer mix (*NOS* terminator and *Cas9*). M: marker, PC: positive control. Analyzed samples are listed in Supplemental Table S3.

Our multiplex PCR strategy employs a three-primer system (Neilan et al. 1997): target-specific forward primers with a poly-dC extension, reverse primers with short tail, and a universal poly-dC primer labeled at the 5′ end with 6-carboxyfluorescein (6-FAM) (Figure 1a). The use of a single FAM-labeled universal primer eliminates the need to individually label each target-specific primer, significantly reducing cost. Moreover, this design allows for the flexible addition of new target-specific primers without requiring additional fluorescent labeling. To control amplification dynamics, the forward primers are used at one-tenth the concentration of the reverse primers. This ensures that forward primers are depleted after several initial cycles, resulting in the incorporation of the poly-dC sequence at the 5′ end of the amplicons. In subsequent cycles, amplification proceeds via the universal poly-dC primer and the target-specific reverse primers, generating fluorescently labeled, poly-dC–tagged fragments. To enable multiplex PCR and fragment analysis of up to 18 target genes (i.e. 15 transgenes and three positive control genes), we comprehensively optimized fragment sizes, amplification regions, primer concentrations, and PCR conditions. Primer concentration was particularly critical, with up to a 10-fold difference required between the highest and lowest concentrations to achieve balanced amplification (Figure 1b and Supplemental Table S1).

As part of our quality control procedures, we routinely use fDET to detect the presence of transgenes in plant resources upon deposition or donation, as well as prior to distribution to users, regardless of whether the plants are reported as transgenic or non-transgenic (Figure 1c-e). These examinations are essential for verifying that the deposited or donated materials contain the transgenes as described by the depositor. Importantly, fDET also plays a critical role in ensuring that transgenic materials possess the intended transgenes and that non-transgenic materials are free from unintended transgene contamination. To evaluate the sensitivity of the method, we prepared bulk seedling samples by mixing three or one transgenic seeds with 100 non-transgenic wild-type seeds (approximately 3 and 1% contamination). Transgenes were successfully detected in both cases, demonstrating that fDET can reliably identify contamination levels as low as 1% (Figure 1f-h). In some cases, we detect additional or unreported transgenes. During 2023 Sep-2026 Mar, we identified 8.4% (25/297) of deposited materials lacked the intended transgenes or contained unintended transgenes. When such discrepancies arise, we investigate the transgene composition and arrangement in detail, communicate our findings to the depositor, and reach a mutual understanding. Following these procedures, the resources are registered in our catalog (Exp-Plant Catalog: https://plant.rtc.riken.jp/resource/home/index.html) and made available for distribution.

In DET, 10 transgene target genes are amplified across four multiplex PCR reactions: (1) *NPTII, HPT* and *BAR*; (2) *GFP, GUS* and *LUC*; (3) 35S promoter and *NOS* promoter; and (4) *NOS* terminator and *Cas9* (Supplemental Table S2). Each reaction also includes *CBP20* as a positive control to confirm amplification success. Since *CBP20* is an Arabidopsis-specific positive control, other target genes should be used when applying DET to other species. The resulting amplicons are analyzed by fragment size using agarose gel electrophoresis (Figure 1i-m).

Together, fDET and DET provide flexible and efficient solutions for transgene detection in plant research and resource management. By accommodating a range of throughput needs and laboratory settings, these complementary methods support reliable verification of genetic status and contribute to the transparency, reproducibility, and biosafety essential for plant biotechnology and the distribution of genetic resources. Notably, fDET and DET facilitate the identification of discrepancies between reported and actual transgene content, offering a valuable tool for quality control and error detection. Future developments may further expand the target range and automation compatibility of fDET, enhancing its utility for large-scale screening and regulatory compliance.

## Materials and Methods

### Plant materials

Leaves of Arabidopsis seedling were frozen in liquid nitrogen and ground into fine powder using ShakeMaster (Bio Medical Science Inc.). Genomic DNA was extracted using the automated DNA Extraction system GENE PREP STAR (KURABO) or GenCheck DNA Extraction Reagent (FASMAC). In our hands, one fully expanded cotyledon yields a sufficient amount of genomic DNA for fDET analysis. Genomic DNA extracted from plants (RIKEN BRC No. psi00201, psi00228, psi00233, psi00286, pst91000) containing various transgenes were pooled (Supplemental Table S3) and 64 ng of pooled DNA was used as a positive control sample for fDET and DET. For the fDET sensitivity assay, genomic DNA was extracted from bulked seedlings of 100 Col seedlings and either three or one seedling(s) of psi00113.

### fDET

Between 1 and 100 ng of genomic DNA was used for multiplex PCR reactions using the QIAGEN Multiplex PCR Kit (QIAGEN), 0.8 µM universal poly-dC primer labeled with 6-FAM, and 2 µL of the fDETac primer mix in a total reaction volume of 50 µL. The sequences and concentrations of each primer included in the fDETac primer mix are listed in Supplementary Table S1. PCR was performed under the following conditions: 95 °C for 15 min; 35 cycles of 94 °C for 30 s, 60 °C for 90 s, and 72 °C for 60 s; followed by a final extension at 72 °C for 30 min. PCR products were diluted 1:240 with ultrapure water and mixed with Hi-Di Formamide (Applied Biosystems) and GeneScan 500 LIZ Size Standard (Applied Biosystems). Samples were denatured at 95 °C for 5 min, placed on ice for 2 min, and then analyzed on a SeqStudio Genetic Analyzer or SeqStudio 24 Flex Genetic Analyzer (Applied Biosystems) with POP-7 polymer using the FragAnalysis or FragmentAnalysis50_POP-7 run module. Fragment analysis was performed using GeneMapper 6 software (Applied Biosystems).

### DET

Between 1 to 100 ng of genomic DNA was used for multiplex PCR reactions using the QIAGEN Multiplex PCR Kit (QIAGEN), 1 µL of the primer mix in a total reaction volume of 10 µL. The sequences and concentrations of each primer are listed in Supplementary Table 3. PCR was performed under the following conditions: 95 °C for 15 min; 38 cycles of 94 °C for 30 s, 60 °C for 90 s, and 72 °C for 30 s; followed by a final extension at 72 °C for 10 min. PCR products were analyzed with 2% agarose gel.

## Data Availability

The data underlying this article are available in the article and in its online supplementary material.

## Acknowledgements

We thank the depositors and donators of valuable biological resources to the RIKEN BRC for sharing their materials with the community. We also thank the members of Experimental Plant Division at RIKEN BioResource Research Center for their technical support.

## Supplemental Materials

Supplemental Table S1. Primers used for fDET Multiplex PCR.

Supplemental Table S2. Primers used for DET Multiplex PCR.

Supplemental Table S3. Arabidopsis lines used in this study.

